# The Neogene-Quaternary diversification trend in the shaping of modern Caribbean mangroves

**DOI:** 10.1101/2022.11.19.517171

**Authors:** Valentí Rull

## Abstract

This paper analyzes the diversification of the Neotropical mangrove flora from the Miocene to the present, using a fairly comprehensive database of 110 pollen records distributed across the whole Caribbean region. A Neogene-Quaternary diversification trend (NQDT) has been identified, characterized by an increase of 25 genera (~78%) with respect to the 7 already existing Paleogene representatives. Only two genera appeared during the Miocene and the maximum increases were observed in the Pliocene-Quaternary transition and the modern-living record. Half of the true-mangrove genera (*Rhizophora, Pelliciera, Acrostichum*) were already present before the Neogene and the others appeared gradually in the Oligo-Miocene (*Crenea*), the Early-Middle Miocene (*Avicennia*) and the Mio-Pliocene (*Laguncularia*). None of the extant associate mangrove genera were present during the Paleogene and all appeared in the Miocene (23 genera) or the Oligo-Miocene transition (3 genera), being the main responsible for the NQDT, in absolute numbers. No regional extinctions were recorded since the Miocene in the Caribbean mangroves, at the generic level. These observations should be complemented with further high-resolution quantitative studies aimed at finding potential causal relationships with climatic, eustatic and paleogeographical shifts.

## Introduction

Mangrove forests cover a large part of worldwide tropical and subtropical coasts (Fig. 1) and play an essential role in the maintenance of terrestrial and marine biodiversity, the ecological functioning of the land/sea intertidal ecotone, and the continuity of global biogeochemical cycles (Chapman, 1976; Saenger, 2002; Nagelkerken et al., 2008; Tomlinson, 2016; Nizam et al., 2022). Mangroves are among the main blue-carbon ecosystems, with high potential to mitigate climate change because of their high rates of carbon accumulation and the significant carbon stocks in their sediments (Fest et al., 2022). Currently, mangroves are among the world’s most threatened ecosystems due to the increasing deforestation rates (25-30% during the last four decades), which could lead to their disappearance within next century (Duke et al., 2017; Goldberg et al, 2020; Worthington et al., 2020). These ecosystems are also highly sensitive to global-change-driven threats such as warming, sea-level rise, increasing extreme meteorological events or invasion by alien species, among others (Gilman et al., 2008; Spalding et al., 2014; Biswas & Biswas, 2019; Wang & Gu, 2021). Knowing past ecological and evolutionary trends, along with the main environmental forcings involved, is useful not only to understand how current mangrove ecosystems have been shaped but also to inform conservation and restoration practices (Berger et al., 2008; Bosire et al., 2008).

**Figure 1.**
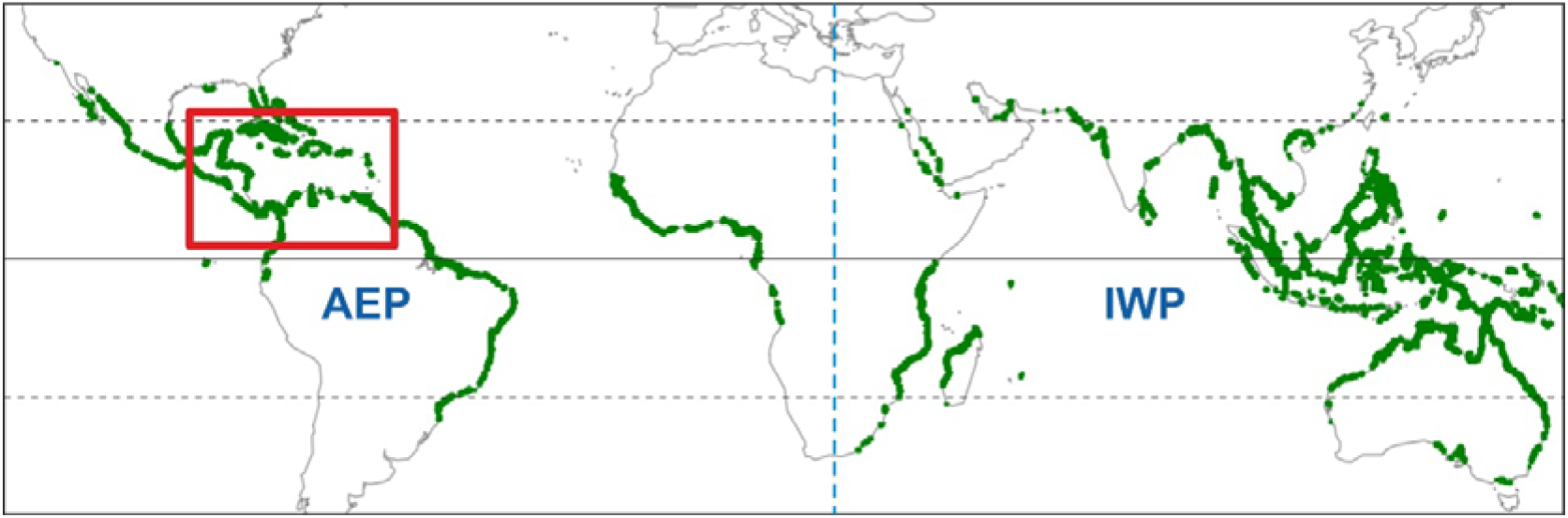
World map of mangroves (green fringes) indicating the Caribbean region (red box). The African continental barrier between the Atlantic-Eastern Pacific (AEP) and the Indo-West Pacific (IWP) biogeographical regions is represented by a dotted blue line in the center. Base map from Spalding et al. (2010).

World mangroves have been subdivided into two major biogeographical domains, the Atlantic-East Pacific (AEP) region and the Indo-West Pacific (IWP) region, separated by the African continental barrier (Fig. 1). The IWP region, with 17 families, 24 genera and 54 species characteristic of mangroves is more diverse than the AEP region, which holds 9 families, 11 genera and 17 species typical of mangrove communities (Duke, 2017). This global biodiversity pattern has led to a variety of hypotheses about the origin and evolution of mangrove communities and the potential environmental drivers involved (Ellison et al., 1999). The Caribbean region has been considered the cradle of Neotropical mangroves, which originated in the Eocene and were dominated by the parent plant of *Lanagiopollis crassa (=Psilatricolporites crassus*), the fossil representative of the extant *Pelliciera rhizophorae* (Rull, 2022a). Other elements of these mangrove communities were the palm *Nypa*, represented by *Spinizonocolpites baculatus/spinosus*, now entinct in the AEP region, and the fern *Acrosticum*, represented by *Deltoidospora adriennis*. The Caribbean mangroves underwent an evolutionary revolution in the Eocene-Oligocene transition (EOT), which has been linked to the abrupt cooling and sea-level drop that characterized this geological boundary. *Pelliciera* was replaced by *Rhizophora* (represented by *Zonocostites ramonae*) as the dominant Neotropical mangrove-forming tree, and this initiated the trend toward the shaping of modern Caribbean mangroves, which are still dominated by this genus (Rull, 2022b). *Pelliciera* did not disappear but became a minor mangrove element that experienced successive Neogene expansion-contraction cycles, until the attainment of its present relict distribution area around a small Central American patch (Rull, 2022c).

The Neogene expansion of modern-like *Rhizophora* mangroves and the reduction of *Pelliciera* was accompanied by a progressive diversification trend that lasted until the Quaternary and led to the current composition of Caribbean mangroves. During the Quaternary, these communities responded to glacial/interglacial climatic and eustatic fluctuations by changing their abundance, composition and distribution patterns. A new element, human disturbance, was introduced in the Holocene that significantly affected mangrove ecology and biogeography, thus contributing to the shaping of current mangrove communities (Rull, 2022d). The Neogene-Quaternary diversification trend (NQDT) was first noted by Graham (1995), who suggested that the Caribbean/Gulf of Mexico mangroves experience a maintained increase in diversity from the Eocene mangroves, characterized by the three basic elements mentioned above, to the current ~27 extant genera. This author also considered the extinct *Brevitricolpites variabilis*, of unknown botanical affinity, as a fourth potential Eocene mangrove component, but further studies have seriously questioned the taxonomic identity and the environmental preferences of the parent plant of this fossil species (Jaramillo & Dilcher, 2001).

The first characterization of the NQDT was based on fossil pollen evidence from some previously published range charts (Germeraad et al., 1968; Muller, 1980; Lorente, 1986), with the addition of a dozen Miocene, Pliocene and Quaternary records (Graham, 1995). In this paper, the NQDT is studied using a newly assembled, updated and fairly complete database retrieved from the original references of 110 Neogene, Quaternary and modern pollen records across the whole Caribbean region. This analysis also benefits from the most updated review of extant mangrove botany (Tomlinson, 2016), which distinguishes true mangrove species from associate elements. The main aim is to provide a robust regional characterization of the NQDT – and, therefore, of the shaping of modern Caribbean mangroves – based on all the available published records.

## Study area

Geographically, the Caribbean region sensu lato is considered, including the Caribbean, Atlantic and Pacific coasts of northern South America, Central America and the Greater/Lesser Antilles, between approximately 5-25 °N and 60-90 °W (Fig. 2). As part of the Neotropical mangroves, the Caribbean representatives are characterized by three main genera of trees: *Rhizophora* (Rhizophoraceae), *Avicennia* (Acanthaceae) and *Laguncularia* (Combretaceae) (Fig. 3). *Rhizophora* is the most abundant and widespread genus and has two Caribbean species (*R. mangle, R. racemosa*), whereas *Avicennia* is represented in the region by three species: *A. germinans, A. bicolor* and *A. shaueriana*. These two genera are of pantropical distribution but their Caribbean species are restricted to the AEP region. The genus *Laguncularía* is monotypic (*L. racemosa*) and exclusive to the AEP region.

**Figure 2.**
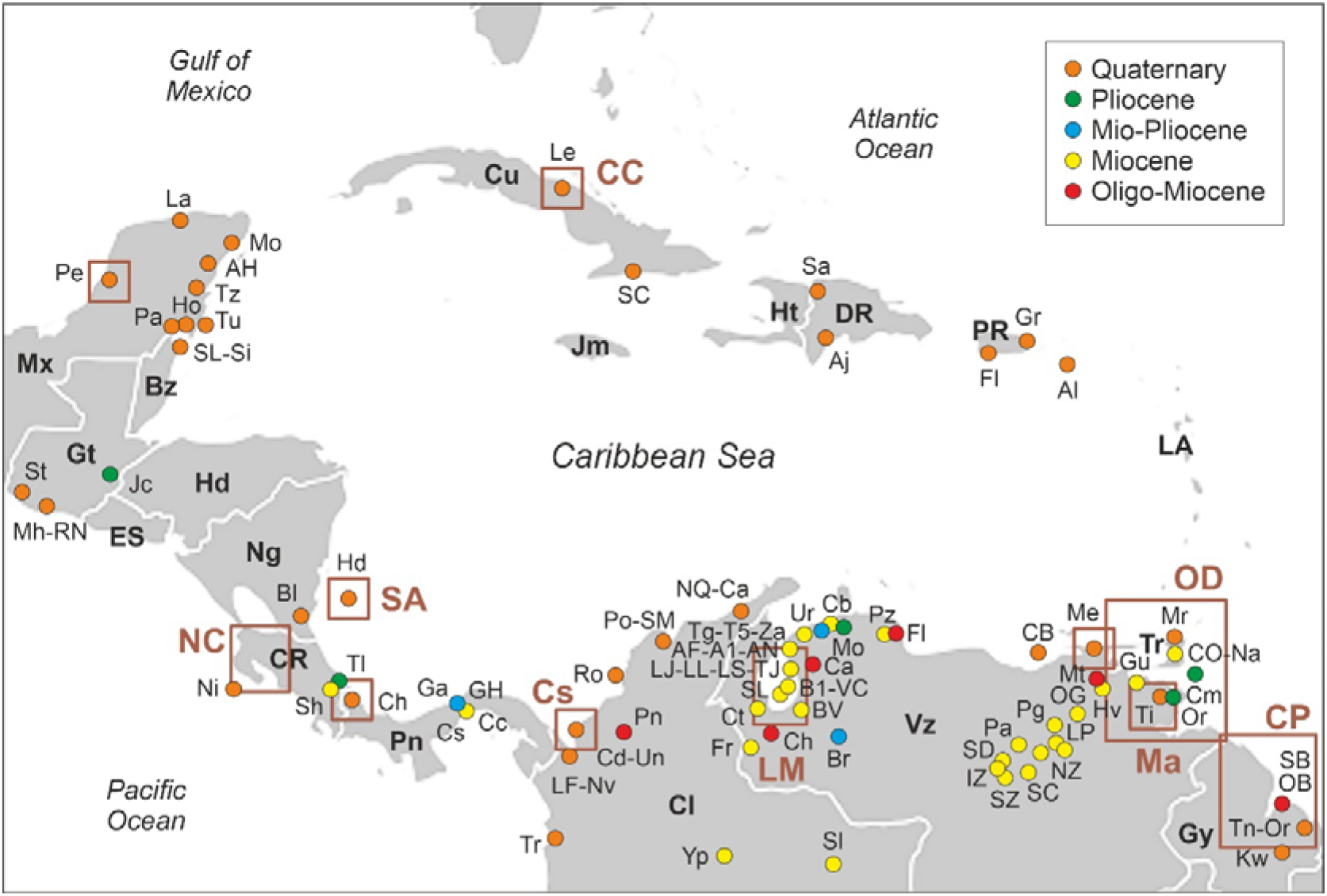
Sketch-map of the Caribbean region indicating the localities with Neogene and Quaternary pollen records (colored dots), and the areas with modern-analog studies (brown boxes). Locality codes according to Table S1. Countries: Bz, Belize, Cl, Colombia; CR, Costa Rica; Cu, Cuba; DR, Dominican Republic; Gt, Guatemala; Gy, Guyana; Ht, Haiti; Hd, Honduras; LA, Lesser Antilles (diverse countries and overseas colonies); Mx, Mexico; Ng, Nicaragua; Pn, Panama; SC, Saint Croix (USA); Tr, Trinidad & Tobago; Vz, Venezuela.

The species of these three genera are called major mangrove elements or true mangroves. To be considered a true mangrove species, the following conditions must be fulfilled (Tomlinson, 2016): 1) occurring only in mangroves and do not extend into terrestrial communities (fidelity); 2) playing a major role in the structure of the community and being able to form pure stands (mangrove—forming trees); 3) possessing special morphological adaptations, as for example aerial roots for gas exchange (pneumatophores) and viviparous embryos; 4) bearing physiological mechanisms for salt exclusion so that they can grow in seawater; and 5) being systematically isolated from their terrestrial relatives, often at the subfamily or family level. Minor true mangrove elements are also restricted and adapted to these communities and are taxonomically isolated at the generic level, but occupy peripheral habitats, rarely form pure stands and are not major structural elements. This is the case of the tree *Pelliciera rhizophorae* (Tetrameristaceae) and the ferns *Acrostichum aureum* and *A. danaefolium* (Pteridaceae) (Fig. 3). *Crenea patentiformis* (Lythraceae) could be considered as a true mangrove species but it is a herbaceous element without the corresponding morphological adaptations.

**Figure 3.**
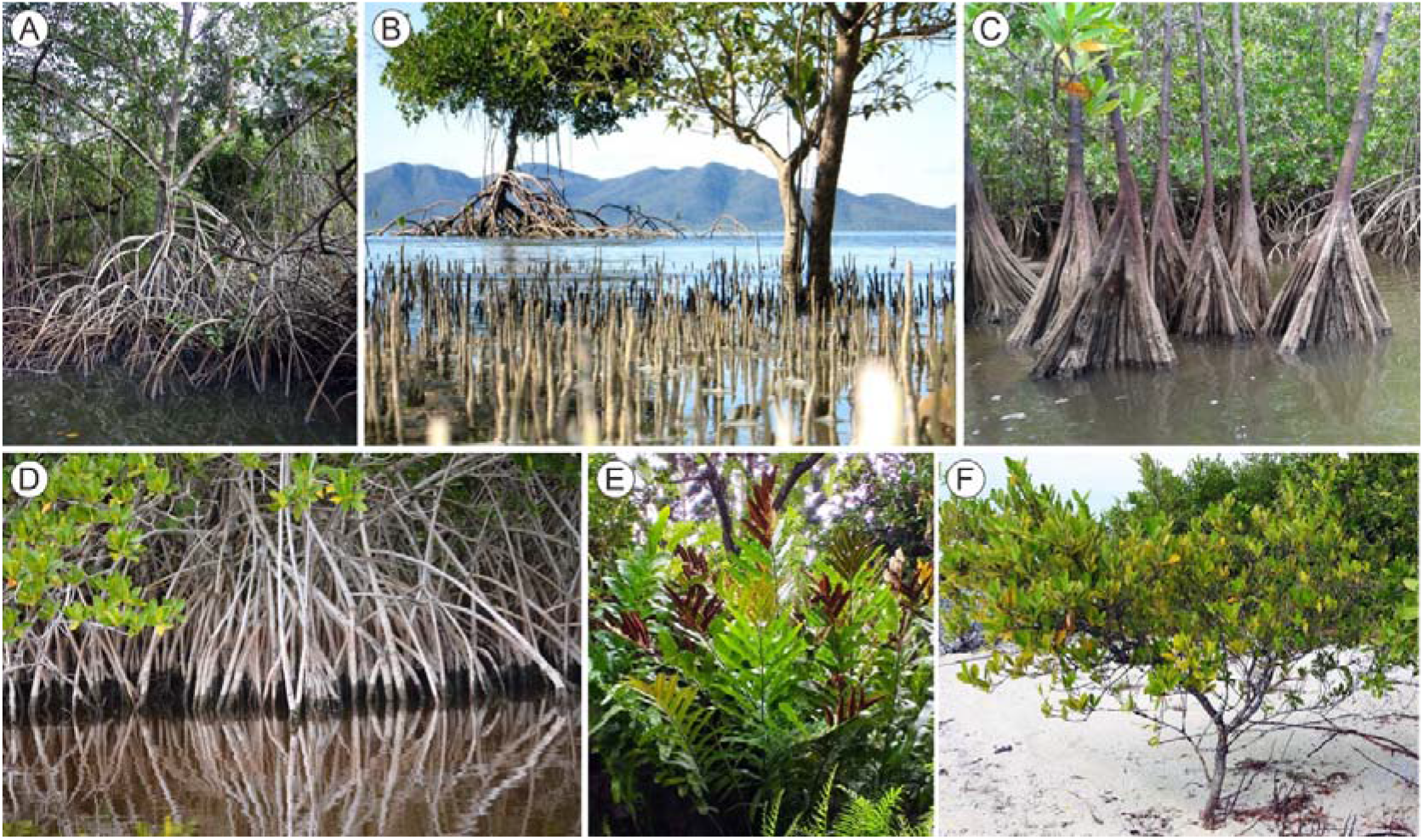
True mangrove species from the Caribbean region. A) *Rhizophora mangle;* B) *Avicennia germinans;* C) *Pelliciera rhizophorae;* D) *Laguncularia racemosa;* E) *Acrostichum aureum;* F) *Conocarpus erectus* (mangrove associate). Reproduced from Rull (2022d).

Other species, known as mangrove associates, can live in mangrove communities but are not restricted to them and can also occur in other habitats, such as coastal swamps, back-mangrove wetlands, salt marshes, riverbanks, beach communities or inland rainforests (Table 1). In addition to the true and associate mangrove flora, a comprehensive floristic study of Neotropical mangroves recognized more than 100 accompanying species, which were useful to define 30 phytosolociological mangrove associations, with 85 of these species as diagnostic – that is, restricted to either one or another of these associations – species (García-Fuentes et al., 2020). The Neotropical mangrove forests show a characteristic sea-inland zonal pattern with no herbaceous understory, characterized by the sequence *Rhizophora-(Pelliciera)- Avicennia-Laguncularia* in the more saline zone dominated by normal tides, *Acrostichum* and *Conocarpus* in the brackish-water back-mangrove swamps and elevated areas under the influence of spring tides, and *Mauritia* and other palms in the more inland freshwater swamps, which mark the transition to the interior savannas and rainforests. Mangrove zonation is influenced by a diversity of biotic and abiotic factors, notably geomorphology, inundation frequency/intensity, salinity, propagule sorting and competition (Lugo & Snedaker, 1974; Rabinowitz, 1978; Smith, 1992; Rull, 1998; Tomlinson, 2016).

**Table 1.** True (in bold) and associate Neotropical mangrove elements, excluding hybrids. Habitats: BC, beach communities; BM, back mangrove; BC, beach communities; CC, coastal communities; CS, coastal swamps; MF, mangrove fringe; RB, river banks; SM, Salt marshes; W, wetlands. Summarized from Tomlinson (2016).

| Genus | Family | Mangrove species | Life form | Habitat |
| --- | --- | --- | --- | --- |
| **Acrostichum** | **Pteridaceae** | **A. aureum, A. danaefolium\*** | **Fern** | **BM** |
| *Amoora* | Meliaceae | *A. cucullata* | Tree | BM |
| *Amphitecna* | Bignoniaceae | *A. latifolia* | Tree | BM/RB |
| *Anemopaegna* | Bignoniaceae | *A. chrysoleucum* | Vine | BM/RB |
| **Avicennia** | **Acanthaceae** | **A. germinans\*, A. bicolor\*, A. shaueriana\*** | **Tree** | **MF** |
| *Batis* | Batidaceae | *B. maritima* | Shrub | SM |
| *Caesalpinia* | Fabaceae | *C. bonduc* | Vine | BM |
| *Conocarpus* | Combretaceae | *C. erectus\** | Tree | BM/W |
| **Crenea** | **Lythraceae** | **C. patentinervis\*** | **Herb** | **MF** |
| *Dalbergia* | Fabaceae | *D. ecastophyllum, D. amerimnion* | Tree/Shrub | BM |
| *Hibiscus* | Malvaceae | *H. tiliaceum* | Tree | BC/CC |
| *Hippomane* | Euphorbiaceae | *H. mancinella* | Tree | BC |
| **Laguncularia** | **Combretaceae** | **L. racemosa\*** | **Tree** | **MF** |
| *Mora* | Fabaceae | *M. oleifera\** | Tree | BM |
| *Muellera* | Fabaceae | *M. moniliformis\** | Tree | RB |
| *Pachira* | Bombacaceae | *P. aquatica* | Tree | BM |
| *Pavonia* | Malvaceae | *P. paludicola\*, P. rhizophorae\** | Shrub | BM/CC |
| **Pelliciera** | **Tetrameristaceae** | **P. rhizophorae\*** | **Tree** | **MF** |
| *Phryganocydia* | Bignoniaceae | *P. phellosperma* | Vine | CS |
| *Pluchea* | Asteraceae | *P. odorata* | Herb | W |
| *Rhabdadenia* | Apocynaceae | *R. biflora* | Vine | BM |
| **Rhizophora** | **Rhizophoraceae** | **R. mangle\*, R. racemosa\*** | **Tree** | **MF** |
| *Rustia* | Rubiaceae | *R. occidentalis* | Tree/Shrub | BM/CS |
| *Scaevola* | Goodeniaceae | *S. plumieri* | Shrub | BC |
| *Tabebuia* | Bignoniaceae | *T. palustris\** | Tree | BM |
| *Thespesia* | Malvaceae | *T. populnea, T. populneiodes* | Tree | BC/CC |
| *Tuberostylis* | Asteraceae | *T. axillaris, T. rhizophorae* | Shrub | BM |

\*Species used by Duke (2017) to characterize the AEP mangroves.

## Methods

This paper analyzes the Neogene-to-present mangrove pollen records available for the Caribbean region. The analysis is based on the sites with published records and the taxa, usually genera, with fossil representatives. Raw data have been retrieved and updated from three previous Caribbean-wide Neogene, Quaternary and modern studies. The original Neogene database was gathered by Rull (2022b, c), and has been complemented here with a number of additional records from the literature, which conforms a dataset of 48 localities (Table S1 of the Supplementary Material). The Quaternary (50 Localities) and modern (12 localities/areas) databases have been taken and updated from Rull (2022d). Modern mangrove standards have been set using patterns of surface pollen sedimentation and their living parent plants. From a taxonomic perspective, not all extant Caribbean mangrove taxa have fossil representatives (Table 3). Besides the already mentioned pre-Neogene records of *Nypa* (already extinct in the AEP region), *Acrostichum, Pelliciera* and *Rhizophora*, others appeared for the first time in the Neogene or the Quaternary and will be the main target of this paper.

**Table 2.** Neogene fossil pollen representatives of extant mangrove genera.

| Genera | Family | Fossil pollen | Reference |
| --- | --- | --- | --- |
| *Acacia\** | Fabaceae | *Acacia/Polyadopollenites mariae* | Graham (1976); Dueñas (1980) |
| *Acrostichum* | Pteridaceae | *Deltoidospora adriennis* | Frederiksen (1985) |
| *Avicennia* | Acanthaceae | *Retitricolpites* sp./*Avicennia* | Lorente (1986); Muller et al. (1987) |
| *Crenea* | Lythraceae | *Verrutricolporites rotundiporus* | Germeraad et al. (1968) |
| *Hibiscus* | Malvaceae | *Echiperiporites estelae* | Germeraad et al. (1968) |
| *Laguncularia* | Combretaceae | *Laguncularia* | Graham (1976) |
| *Pachira* | Bombacaceae | *Bombacacidites baculatus* | Lorente (1986) |
| *Pelliciera* | Pellicieraceae | *Psilatricolpoites crassus* | Wijmstra (1968) |
| *Rhizophora* | Rhizophoraceae | *Zonocostites ramonae* | Germeraad et al. (1968) |

\*Not included in Table 1 but considered to be a mangrove element in the cited references.

**Table 3.** First Neogene appearances and ranges of occurrence of pollen from mangrove genera with Neogene fossils (Table 2), by localities. Site abbreviations in Table S1. *Pel, Pelliciera; Rhi, Rhizophora; Cre, Crenea; Hib, Hibiscus; Acr, Acrostichum; Pac, Pachira; Aca, Acacia; Avi, Avicennia; Lag, Laguncularia*. E, Early; M, Middle; L, Late.

| Age | *Pel* | *Rhi* | *Cre* | *Hib* | *Acr* | *Pac* | *Aca* * | *Avi* | *Lag* | Sites |
| --- | --- | --- | --- | --- | --- | --- | --- | --- | --- | --- |
| Pliocene | X | X |  | X | X |  | X | X | X | Or/Cm/Tl/Ga/Jc/SB |
| L Mio-Pliocene | X | X | X | X | X | X | X |  |  | AF/Br/Hv |
| L Miocene | X | X | X | X | X | X | X |  |  | A1/Za/Gu/SL/T5 |
| M Mio-Plio | X | X | X | X | X | X | X |  |  | Mo/CO |
| M-L Miocene | X | X | X | X | X | X |  |  |  | VC/Yp/BV |
| E Mio-Pliocene | X | X | X | X | X | X |  |  |  | Ct/B1 |
| E-L Miocene | X | X | X | X | X | X |  |  |  | Tg/Ur/Cb/OG |
| E-M Miocene | X | X | X | X | X | X | X |  |  | NZ/Pa/Pg/LP/Pz/LL/LS/LJ/Ch |
| E Miocene | X | X | X | X | X | X | X |  |  | TJ/SC/SD/Sh/Sl/SZ/IZ/Cc/Cs/GH/AN |
| L Oligo-E Mio | X | X | X | X | X | X | X |  |  | Mt/Pn |
| Oligo-E Mio | X | X | X | X | X |  |  |  |  | Ca/Fr |
| Oligo-Miocene | X | X | X | X |  |  |  |  |  | Fl/SB |

\*Not included in Table 1 but considered to be mangrove elements in the corresponding records (Table S1).

Only genera have been considered, due to the unfeasibility of distinguishing among species, on the basis of solely pollen morphology. Chronologically, the latest version of the International Chronostratigraphic Chart is adopted, where the Neogene ranges from 23.03 Ma to 2.58 Ma, with the Mio-Pliocene boundary situated at 5.33 Ma (Cohen et al (2013). In this framework, the Quaternary began at 2.58 Ma and the Holocene encompasses the last 11.7 kyr. Emphasis is placed on diversification, or net diversity increase, understood as the balance between speciation and extinction (Cracraft, 1985), and is expressed in terms of taxa richness.

## Results

### Neogene

Table 4 displays the first appearances and the ranges of occurrence of Caribbean genera with Neogene fossil pollen representatives, according to Table 3, indicating the localities from where the evidence has been retrieved. In the case of *Pelliciera, Rhizophora* and *Acrostichum*, the first appearances were recorded before the Neogene, in the Early Eocene, Middle Eocene and Late Cretaceous, respectively, as documented in the corresponding literature (Rull, 2022a). The first appearances of *Hibiscus* and *Crenea* were recorded in Oligo-Miocene sediments from Venezuela (Falcón) (Rull & Poumot, 1997) and Guyana (Shelter Belt) (Van der Hammen & Wijmstra, 1964), respectively. *Pachira* and *Acacia* appeared for the first time in the Late Oligocene-Early Miocene of Venezuela (Maturín) (Helenes & Cabrera, 2002) and Colombia (Planeta Rica) (Dueñas, 1980), respectively. *Avicennia* was first recorded in the Early-Middle Miocene of Venezuela (Los Pobres-1) (Lorente, 1986), whereas *Laguncularia* appeared for the first time probably in the Late Pliocene of Costa Rica (Talamanca) (Graham & Dilcher, 1998). This first *Laguncularia* record is based on the presence of a pollen type called Unknown-3 in the original reference (Graham & Dilcher, 1998) and identified here as likely *Laguncularia*, on the basis of the images provided in it. A slightly earlier record is available for the Gulf of Mexico, in a locality that is outside the study area of this paper but not far from the Yucatan Peninsula, where pollen identified as *Laguncularia* appeared in the Late Miocene (Graham, 1976). According to these records, six of the nine extant mangrove genera (if we consider *Acacia* as a mangrove element) with known fossil pollen emerged during the Neogene. With the Miocene and Pliocene appearances of *Avicennia* and *Laguncularia*, respectively, all true Neotropical mangrove elements were already present in the Caribbean region by the end of the Neogene.

**Table 4.** First Quaternary appearances and ranges of occurrence of pollen from mangrove genera with Quaternary representatives, by localities. *Cas, Cassipourea; Pte, Pterocarpus; Ran, Randia; Fic, Ficus, Con, Conocarpus; Bat, Batis; Hip, Hippomane; Tab, Tabebuia; Cae, Caesalpinia; Dal, Dalbergia*. E, Early; M, Middle; L, Late.

| Age (cal kyr BP) | *Cas* * | *Pte* * | *Ran* * | *Fic* * | *Con* | *Bat* | *Hip* | *Tab* | *Cae* | *Dal* | Sites |
| --- | --- | --- | --- | --- | --- | --- | --- | --- | --- | --- | --- |
| L Holocene (1-0) |  |  |  |  | X |  |  |  |  |  | Nv/Ro/LF |
| L Holocene (2-1) |  |  | X | X | X | X | X | X | X | X | Hd/Ca/Po |
| L Holocene (3-2) |  |  |  | X | X |  |  |  |  |  | Mo |
| L Holocene (4.2-3) |  | X |  | X | X |  |  |  |  |  | Gr/La/Na/SC/Pe |
| M Holocene (5-4.2) |  |  |  | X | X |  |  |  |  |  | Pa/Ga |
| M Holocene (6-5) |  |  |  | X | X |  |  |  | X |  | Ho/Ni/To/En/Le |
| M Holocene (7-6) | X |  |  | X |  | X | X | X |  |  | Ti/NQ/Mr |
| M Holocene (8.2-7) |  |  | X | X | X | X |  |  |  |  | Le/Tz |
| E Holocene (>8.2) | X | X | X |  |  |  |  |  |  |  | GA |

\*Not included in Table 1 but considered to be mangrove elements in the corresponding records (Table S1).

### Quaternary

Quaternary records are mostly based on Holocene sequences, as Pleistocene records are absent, except for a few Late Pleistocene sections (Rull, 2022d). The oldest published records correspond to the last glacial cycle (~130-70 cal kyr BP), when mangrove pollen assemblages were dominated by *Rhizophora, Avicennia* and *Acrostichum* (Van der Hammen, 1963; González & Dupont, 2009; González et al. 2008). After the last glaciation, Early Holocene sequences are mostly absent from the Caribbean region and the records restart in the Middle Holocene (8.2 cal kyr BP onward) (Rull, 2022d). According to the available evidence, three genera (*Cassipourea, Pterocarpus* and *Randia*) were present in Central America (Panama) since the beginning of the Holocene (Bartlett & Barghorn, 1973). Other three genera (*Ficus, Conocarpus* and *Batis*) appeared in Cuba (La Leche) and Mexico (Lake Tzib) at the beginning of the Middle Holocene (Peros et al., 2007; Carrillo-Bastos et al., 2010). The first appearances of *Hippomane* and *Tabebuia* were recorded in Trinidad (Maracas) and Venezuela (Tigre), respectively, during the Middle Holocene (Ramcharan & McAndrews, 2006; Montoya et al., 2019). *Caesalpinia* also appeared during the Middle Holocene in Mexico (Rio Hondo) and *Dalbergia* was first recorded in the Late Holocene of Colombia (Honda) (González et al., 2010; Aragón-Moreno et al., 2018). However, due to the above-mentioned absence of pollen records for most of the Pleistocene, it is not possible to know whether these genera were younger than Holocene.

### Modern

The available studies on modern pollen deposition add two genera to the list of sedimentary pollen: *Rhabdadenia*, from the Orinoco delta and surrounding areas (Rull & Vegas-Vilarrúbia, 1999; Hofman, 2002) and *Scaevola*, from Cuba (Cayo Coco) (Davidson, 2007). The remaining genera of Table 1 (*Amoora, Amphitecna, Anemopaegna, Mora, Muellera, Pavonia, Phryganocydia, Pluchea, Rustia, Thespesia and Tuberostylis*) are only known by their living species.

### Conclusions and discussion

Fig. 4 is a graphical synthesis of the outcomes of this analysis, from which the following points may be highlighted:

- A total of 32 genera have been identified in the Neogene, Quaternary and modern mangrove record, of which six are true mangroves and the remaining 26 are associates (21) or potential associates (5).
- Overall, the NQDT was characterized by an increase of 25 genera (~78%) with respect to the 7 already existing Paleogene representatives.
- Only two genera appeared during the Miocene and the maximum increases were observed in the Pliocene-Quaternary transition (PQ; 10 appearances) and the modern-living record (ML; 11 appearances).
- Half of the true-mangrove genera (*Rhizophora, Pelliciera, Acrostichum*) were already present before the Neogene and the others appeared gradually in the Oligo-Miocene (OM) (*Crenea*), the Early-Middle Miocene (*Avicennia*) and the Mio-Pliocene (MP) (*Laguncularia*).
- None of the extant associate mangrove genera were present during the Paleogene and all appeared in the Miocene (23 genera) or the OM transition (3 genera), being the main responsible for the NQDT, in absolute numbers.
- No regional extinctions were recorded since the Miocene in the Caribbean mangroves, at the generic level.

**Figure 4.**
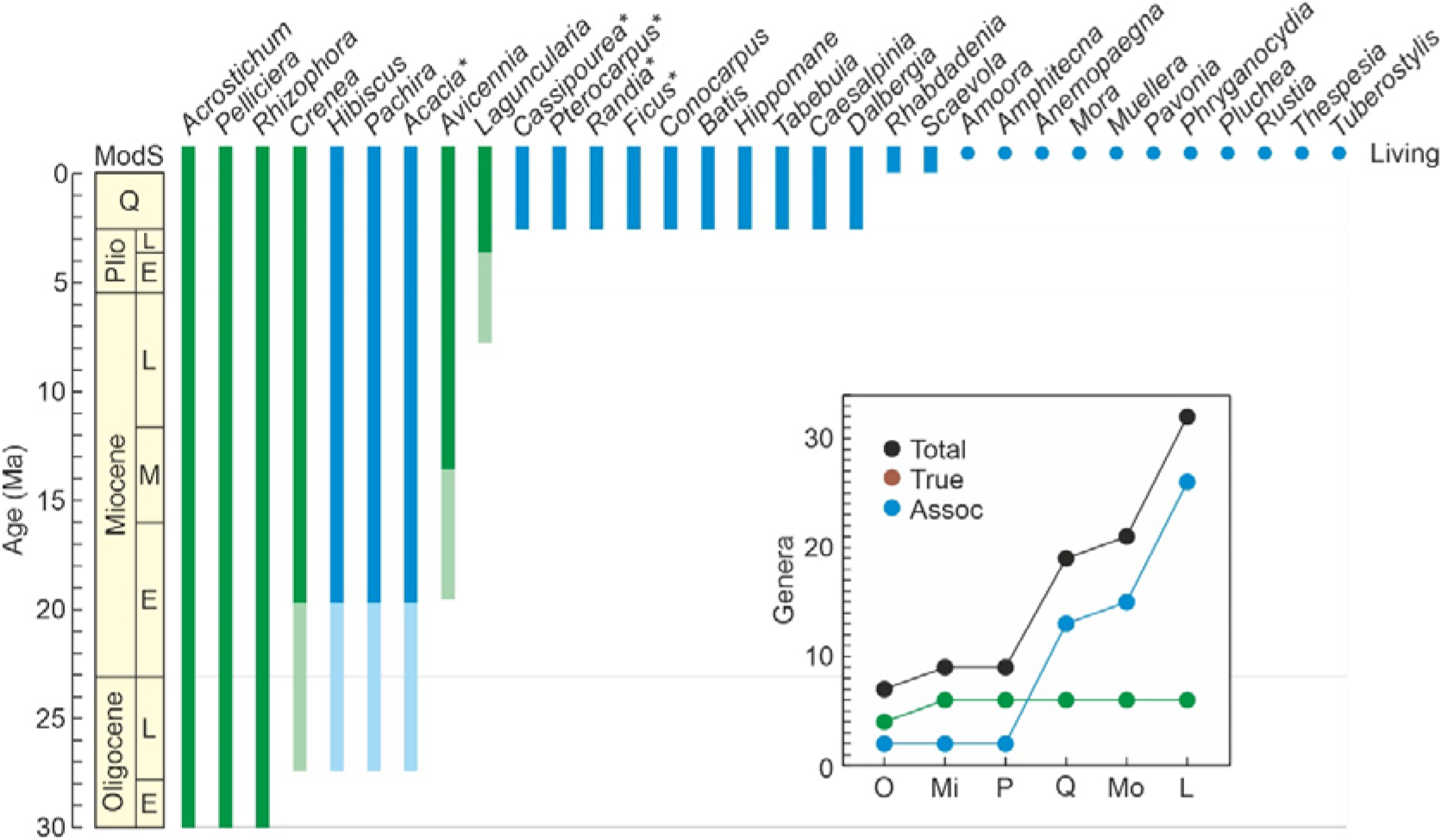
Synthetic range chart of the extant genera from the Caribbean region and their Neogene, Quaternary and modern pollen representatives. Bars are pollen ranges and dots represent living species. True mangrove genera are in red and associates in blue (*genera not listed in Table 1 but considered as mangrove associates in the corresponding records). Chronological uncertainty in the first appearances are indicated by light green/blue bars. Gray bands in the richness plot represent phases of diversification acceleration. 0, Oligocene; Mi, Miocene; P, Pliocene; Mo, modern; L, living.

Some of these points deserve specific comments. The PQ increase occurred mostly during the Pleistocene and Early Holocene, for which no records are available, except for a few late Pleistocene sequences. Therefore, this apparent diversification acceleration emerges mostly from the comparison of pollen records from Pliocene rocks and unconsolidated Mid-Late Holocene sediments, with contrasting taphonomic conditions, especially in relation to pollen preservation. To bridge this gap, it would be essential to study Pleistocene sediments, which are present mainly in marine cores, as for example those retrieved in the Venezuelan Cariaco Basin (site CB), which encompass long Pleistocene sequences (Haug et al., 1998; Yarincik & Murray, 2000). Regarding the ML diversity increase, the presence of methodological artifacts could not be dismissed, as modern records are based on pollen from surface sediments, whereas living records refer to the occurrence of parent plants in the extant flora. These records are hardly comparable, as the living plant record is most likely complete, whereas the modern record may be more fragmentary and affected by idiosyncratic features related with pollen production, dispersal, sedimentation, preservation, reworking and other taphonomic processes. Ideally, fossil pollen records should be compared with studies on modern pollen sedimentation, rather than with living taxa.

Other interesting observations are that the main structural mangrove elements are far less diverse than associated taxa and attained their present richness during the Miocene, before the diversification of associate taxa. This indicates that the structural features of Caribbean mangrove communities were already set at the beginning of the Neogene and the responsible taxa (mainly *Rhizophora* and *Avicennica*) were successful enough to remain until today, thus providing the structural and functional conditions for the survival and diversification of the associate flora. The absence of extinctions and community turnovers supports the evolutionary resilience of Neogene Caribbean mangroves, whose main ecological features would have established in the Miocene and remained until today, despite the occurrence of significant climatic, eustatic and paleogeographical shifts (Westerhold et al., 2020). Establishing potential causal relationships between Neogene mangrove diversification and external environmental shifts is still premature due to the comparatively low chronological resolution and the scarcity of quantitative pollen records. However, it is noteworthy that the main diversification increase documented to date, the PQ acceleration, occurred during the Pleistocene, characterized by intense and recurrent climatic and eustatic shifts linked to the glacial-interglacial cycles. This coincidence deserves further consideration and the development of studies aimed specifically at analyzing potential causal relationships.

The general diversification patterns recorded in this analysis are consistent with previous observations by Graham (1995) and provide additional and complementary information based on a fairly comprehensive, in geographical and chronological terms, database encompassing the whole Caribbean region. In addition to provide direct evidence on mangrove evolution, these pollen records are able to furnish calibration points to improve chronological interpretations in molecular phylogenetic and phylogeographic studies of mangrove taxa (Graham, 2006; Triest, 2008; Lo et al., 2014; Xu et al., 2017; Takayama et al., 2021).

## Acknowledgements

No funding was received specifically for the development of this research.

